# The self-organization of plant microtubules in three dimensions enable stable cortical localization and sensitivity to external cues

**DOI:** 10.1101/210138

**Authors:** Vincent Mirabet, Pawel Krupinski, Olivier Hamant, Elliot M Meyerowitz, Henrik Jönsson, Arezki Boudaoud

**Affiliations:** Reproduction et Développement des Plantes, Univ. Lyon, ENS de Lyon, UCB Lyon 1, CNRS, INRA,F-69364 Lyon, France.; Computational Biology and Biological Physics Group, Department of Theoretical Physics, Lund University, S-221 00 Lund, Sweden.; Sainsbury Laboratory, University of Cambridge, Bateman Street, Cambridge CB2 1LR, UK.; Division of Biology and Biological Engineering, California Institute of Technology, Pasadena, CA 91125, USA.; Howard Hughes Medical Institute, California Institute of Technology, Pasadena, CA 91125, USA.

## Abstract

Many cell functions rely on the ability of microtubules to self-organize as complex networks. In plants, cortical microtubules are essential to determine cell shape as they guide the deposition of cellulose microfibrils, and thus control mechanical anisotropy in the cell wall. Here we analyze how, in turn, cell shape may influence microtubule behavior. Using a computational model of microtubules enclosed in a three-dimensional space, We show that the microtubule network has spontaneous configurations that could explain many experimental observations without resorting to specific regulation. In particular, we find that the preferred localization of microtubules at the cortex emerges from directional persistence of the microtubules, combined with their growth mode. We identified microtubule parameters that seem relatively insensitive to cell shape, such as length or number. In contrast, microtubule array anisotropy depends strongly on local curvature of the cell surface and global orientation follows robustly the longest axis of the cell. Lastly, we found that the network is capable of reorienting toward weak external directional cues. Altogether our simulations show that the microtubule network is a good transducer of weak external polarity, while at the same time, it easily reaches stable global configurations.

**Author summary:** Plants exhibit an astonishing diversity in architecture and shape. A key to such diversity is the ability of their cells to coordinate and grow to reach a broad spectrum of sizes and shapes. Cell growth in plants is guided by the microtubule cytoskeleton. Here, we seek to understand how microtubules self-organize close to the cell surface. We build upon previous two-dimensional models and we consider microtubules as lines growing in three dimensions, accounting for interactions between microtubules or between microtubules and the cell surface. We show that microtubule arrays are able to adapt to various cell shapes and to reorient in response to factors such as signals or environment. Altogether, our results help to understand how the microtubule cytoskeleton contributes to the diversity of plant shapes and to how these shapes adapt to environment.

## Introduction

Despite their amazing diversity in shapes, biological organisms share some common structural components at the cellular level. Among those, one of the best conserved proteins across eukaryotes, tubulin, assembles into protofilaments, which in turn form 25 nm nanotubes known as microtubules, usually made of 13 protofilaments. The network of microtubule is highly labile and can reshape itself in a matter of minutes. In plants, microtubules form superstructures before (the preprophase band), during (the spindle) and after (the phragmoplast) cell division. Plant microtubules also form dense and organized arrays at the periphery of the cell during interphase [1] and these arrays are known as cortical microtubules (CMTs). The behavior of CMTs has been studied extensively at the molecular level [2]. One of their main functions is to guide the trajectory of the transmembrane cellulose synthase complex and thus to bias the orientation of cellulose microfibrils in the wall. This in turn impacts the mechanical anisotropy of the cell wall and controls growth direction [3], [4], [5], [6]. This function explains why most mutants impaired in microtubule-associated proteins exhibit strong phenotypic defects [7].

Whereas this provides a clear picture on how microtubules impact cell shape, in turn how cell shape impacts microtubule behavior has been much less explored. There is evidence that shape-derived mechanical stress can bias microtubule orientation towards the direction of maximal tension, both at the tissue and single cell scales [8], [9]. The molecular mechanism behind this response remains unknown. In addition, cell geometry may also affect microtubule behavior, independently of the presence of mechanical stress in cell walls. This is what we explore here. Before doing so, we first review the main features of the microtubule network.

The molecular basis for microtubule dynamics is rather well established. Consistent with the absence of centrosome in land plants, microtubule nucleation is dispersed in plant cells, as it occurs at the cell cortex [10], along existing microtubules during branching events [11], [10], and at the nuclear envelope [12]. As they grow, microtubules form stiff and polar structures. They can alternate growth, pause and shrinking at the so-called plus end [13], whereas they mainly shrink or pause at the minus end [14]. The combination of an average shrinkage at the minus end and dynamic instability at the plus end leads to an overall displacement of the microtubule, also called hybrid treadmilling. This is the main cause for microtubule encounters, and thus for reshaping the microtubule network [13], [14].

When one microtubule encounters another microtubule, different outcomes can be observed: if the encounter angle is shallow, zippering can occur, i.e. the growing microtubule bends and continues its polymerization along the encountered microtubule, which leads to the creation of microtubule bundles; if the encounter angle is steep, crossover can occur, i.e. a microtubule polymerizing without deviating its trajectory and crossing over the encountered microtubule; or alternatively catastrophe is triggered, i.e. a rapid plus end shrinkage after contact with the encountered microtubule.

Such selective pruning of microtubules may explain how microtubules can form parallel arrays from initially random orientations, and conversely change the net orientation of their arrays over time, through a phase of randomisation [15], [16]. The presence or absence of different microtubules associated proteins (MAPs) can modulate the stability of microtubules or their capacity to form bundles and to self-organize. For instance, the microtubule severing protein Katanin accounts for most of the pruning events at crossover sites [17].

The microtubule network is a typical example of a self-organizing system, where properties of individual elements and their interactions induce specific and sometimes counter-intuitive global properties. To predict how regulation at the level of each microtubule can give rise to specific global outcomes, one can resort to computational models. Modeling approaches have been developed, simplifying microtubule interactions by restricting them to the plasma membrane, i.e. a simpler 2D space [18], [19]. In those agent-based models, several microtubule properties were coded and interactions between CMTs, based on these properties, were simulated. The outcome is an emergent network, whose characteristics can be analyzed. For instance, increased microtubule severing was predicted to generate a larger number of free microtubules, more amenable to bundle into aligned arrays [20] and this was observed in experiments [21].

So far, most of the microtubule models have been implemented in a 2D space. A major outcome of such models was to demonstrate that global orientations of the network can spontaneously emerge from the interactions between microtubules. Many combinations of parameters and behaviors have been studied: instability at the plus end [22], [23], role of zippering [20], [24], [25], [26], nucleation modes [26], [27], and severing [28]. Beyond the differences, the fact that a global orientation emerges in many of these combinations suggests that reaching one global orientation is a robust feature of microtubule networks. Conversely, the diverse combinations of microtubule properties provide different scenarios for the fine-tuning of the network structure and stability of this emerging behavior.

Some aspects of cell geometry were related to microtubule behavior in certain simulations. Simulations showed how different directional biases in nucleation can induce an ordering of the array toward directions that are correlated to cell geometry [22] and [29]. Further, branching nucleation rules can elicit handedness of the global direction of microtubule arrays, provided that the branching is biased toward one direction [30]. Other studies used a simulation space where borders, analog to cell edges, induce more or fewer catastrophe events or are more or less permissive toward microtubule growth [31], [22], [30] and [15]. Most studies concluded that a global orientation of microtubules can be correlated to cell face orientations.

The contribution of the third dimension to microtubule behavior has started to be investigated. Computational models for animal systems have focused on 3D considerations but the nucleation hypotheses are too different from that in plants to be transposed directly [32], [33]. Accordingly, fully 3D models suited for plants are still lacking: almost all existing studies have only considered microtubules living on surfaces embedded in 3D [34], [30], [31], [35]. In [22], a 2D model was extended into a full 3D model but it did not include cell boundaries, which yielded microtubules distributed over the whole simulated domain, in contrast with the cortical localization of microtubules *in planta*.

In this paper, we explore the influence of 3D cell shape on the basic properties of a dynamic microtubule network. We do not assume that microtubules live on a 2D surface; rather, we simulate a closed volume where microtubules are more or less free to grow in all directions. Anchoring to the membrane is not imposed by the model and instead becomes a variable in the model. Using this framework, we investigate how microtubule interaction with the membrane influences microtubule dynamics. Our study also addresses the relative contributions of cell shape, microtubule interactions, and external directional cues in network organization.

## Results

### Microtubules become cortical in a 3D space because of their directional persistence and growth mode

We modelled microtubules as a set of line segments that nucleate, grow, shrink, and interact between them and with the cell surface represented as a triangular mesh (Methods). Nucleation occurs randomly at the surface. Growth occurs from the plus end with a small directional noise that is related to the persistence length of microtubules. Shrinkage occurs at the minus end. A microtubule that encounters a previously existing microtubule either changes direction to that of the previous microtubule (if the encounter angle is steep) or undergoes catastrophe (complete shrinkage). We considered two types of interaction with the cell surface: strong anchoring, whereby a microtubule that reaches the surface continues growing tangentially to the surface, and weak anchoring, whereby microtubules are prevented from leaving the cell interior. Typical simulated shapes and simulation runs are shown in Fig 1.

**Fig 1.**
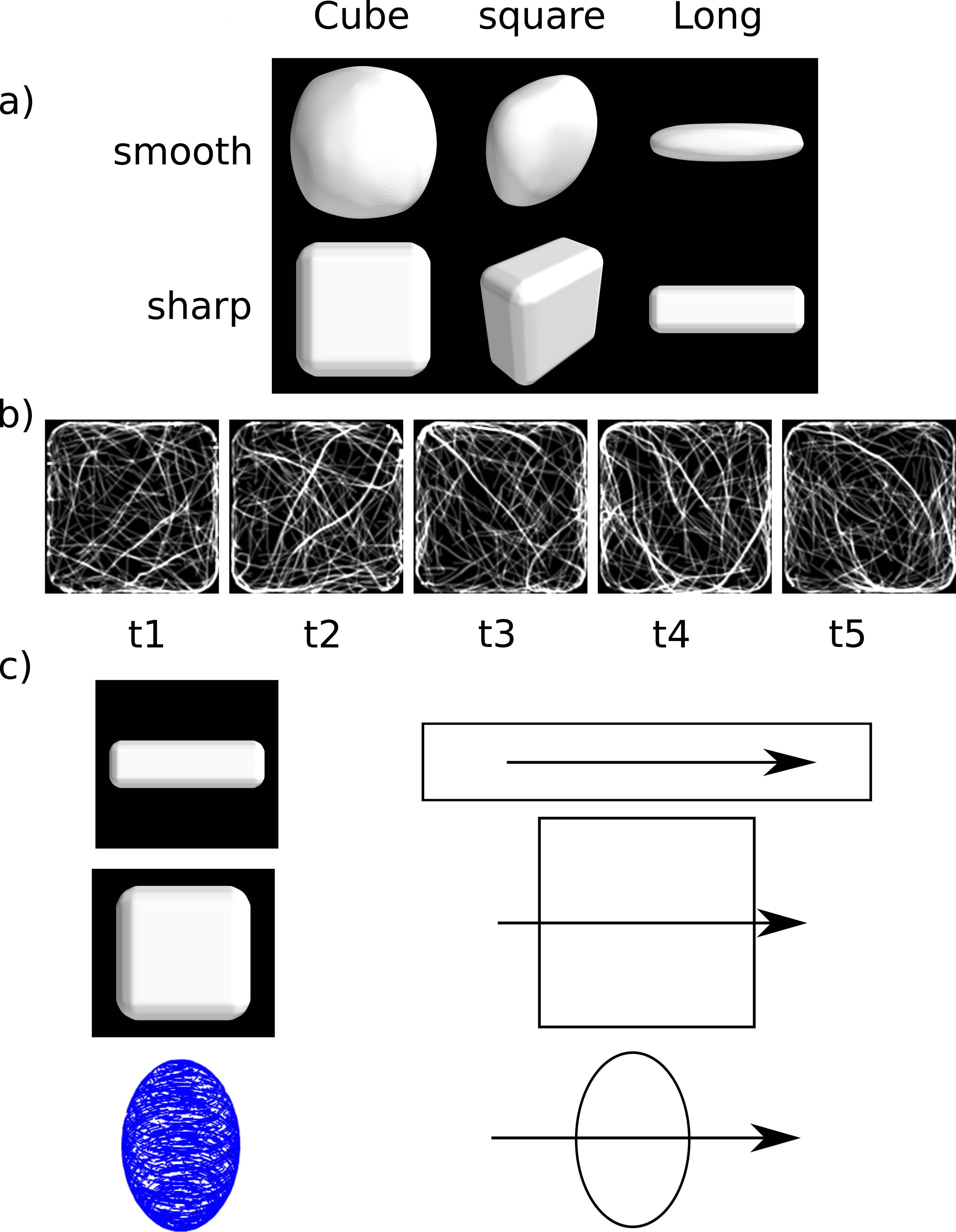
Shapes of simulated cells. (A) Shapes of the envelope of the cell. First row: smooth shapes. Second row: sharp shapes. Scales are not respected; dimensions: sharp cube (8.8µm*8.8µm*4.8µm), smooth cube (9µm*9µm*4.7µm), sharp long (4.8*4.8*15.6µm). (B) Snapshots of a simulation run over 50000 timesteps (corresponding to approximately 100 minutes). Each snapshot is taken after 10000 timesteps (~20 min). The pictures show a z-projection of a treated output (Methods). (C) Reference of angles for the different shapes used here.

We first considered the anchoring of the microtubules to the membrane. There are proteins that have been shown to be associated with both microtubules and a plasma membrane component in plants. For instance, CELLULOSE SYNTHASE INTERACTIVE PROTEIN 1 (CSI1) interacts with both CMTs and the cellulose synthase (CESA) complex [36], [6] and CSI1 was also proposed to stabilize the microtubule network [37].

Yet, the influence of CMT-CESA interactions on the microtubule network is still poorly understood. Thus, we took advantage of the 3D nature of our model to study the impact of the anchoring rule to the membrane on the global parameters of the microtubule network.

In case of the strong anchoring, microtubule growth is biased so as to stay near the membrane, as if anchoring proteins were highly concentrated. As expected, strong interactions led to a surface-localized cortical zone with microtubules trajectories embedded in the plane parallel to the mesh (Fig 2A).

**Fig 2.**
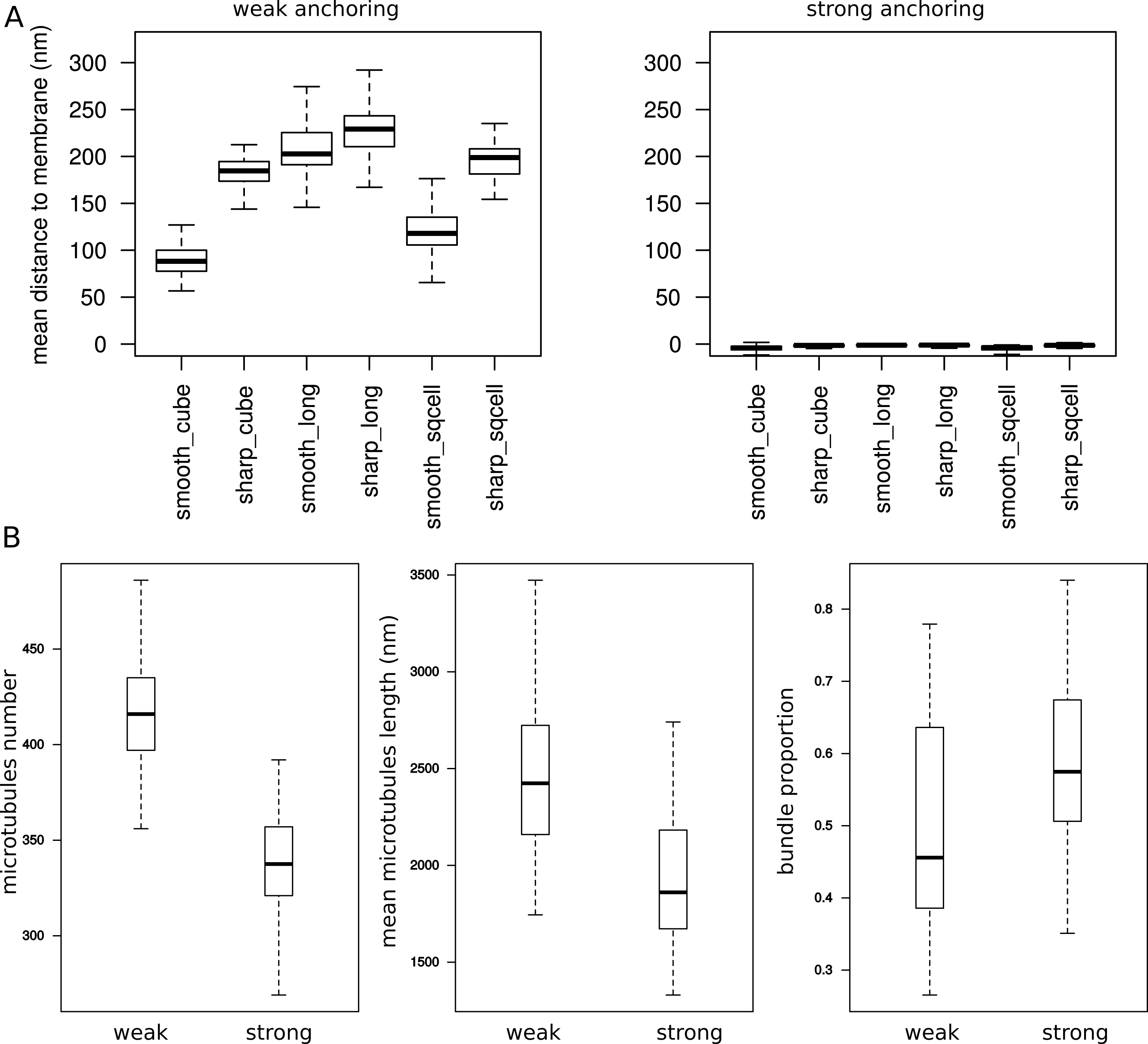
Influence of the anchoring to the membrane. (A) Mean distance of microtubules to the membrane according to cell shape and to weak/strong anchoring to the membrane. (B) Effect of anchoring on three properties of the microtubule network. Left: total number of microtubules. Middle: mean microtubule length. Right: proportion of tubulin in bundles. Simulation parameters (Methods): n_p_ = 1, 2, or 3; n_s_=0.001; r_d_=2.5%; α=40.1°.

We then tested the microtubule response when anchoring to the membrane is weak. In that case, microtubules can crawl in every direction, but sometimes as they encounter the membrane, their direction may be transiently tangent to the membrane. Even if weak anchoring allows microtubules to travel through all the volume of the cell, we find that such weak interaction with the membrane is enough to elicit the existence of cortical microtubules. In those conditions, the predicted distance between microtubules and membrane would be from 50 to 250 nm (Fig 2A). Therefore, the three-dimensional nature of our model helps us demonstrate that strong anchoring is not required for the presence of large populations of cortical microtubules in plant cells: the directional persistence, together microtubule growth mode, can cause such subcellular localization.

As the microtubules tend to stay at close to the membrane, they also bundle, at proportions varying from 30% to 75% (Fig 2B). The ability for the microtubule network to generate a spontaneous bundled structure is consistent with 2D models. Strikingly, this effect is also present in the case of weak anchoring.

### Strong anchoring decreases microtubule number and length, and increases microtubule bundling

Independently of the encounter rule, weak anchoring strength increases the total number and the size of microtubules (by about 20%, in length and in number) when compared to strong anchoring. Weak anchoring strength also generated less bundling (reduction of about 20%). This effect could be due to the fact that a weak anchoring to the membrane allows microtubules to crawl inside the cell, thus diminishing the encounter probability. Consequently, microtubules weakly bound to the membrane have more space to grow and are less subject to blockage or bundling by other microtubules (Fig 2B). Lastly, we found that anchoring strength significantly affected the anisotropy of the microtubule arrays. High anchoring increased the anisotropy of the network by about 10% (Fig 4C).

### The number and length of microtubules, as well as the proportion of microtubules in bundles, are relatively independent of cell shape

Next, we used our model to determine the consequences of changing the cell shape on global properties of the network. We simulated the network in three main sharp shapes represented on Fig 1A. The results from the simulations indicate that these shapes only have a marginal effect on the number of microtubules (less than 10% difference), on the length of microtubules (5 to 15% difference) and on the proportion of bundles (less than 15% difference). Overall, elongated cells have more microtubules, more bundles, and longer microtubules, while cubes show the lower values (Fig 3).

**Fig 3.**
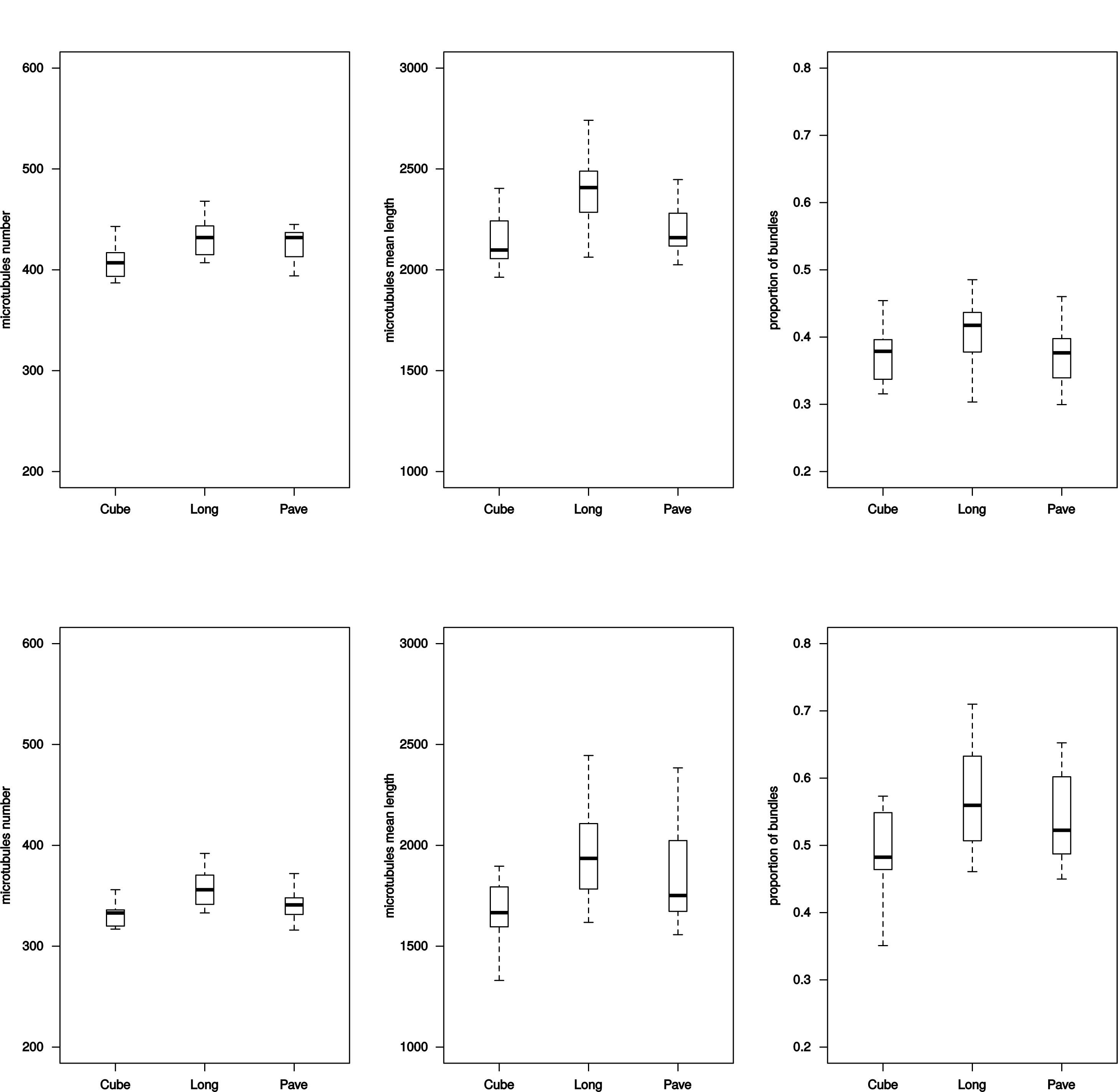
Influence of shape on three global properties of the network. From left to right: number of microtubules, mean microtubule length, and proportion of tubulin in bundles. Top row: weak anchoring; bottom row: strong anchoring. For details on shapes see Fig 1.

### Microtubule array anisotropy is not influenced by the cell global shape but by its curvature

We also analyzed the effect of cell shape on the microtubule array anisotropy, averaged over the cortex (see Methods). As microtubule array anisotropy is skewed towards low values (found inside the cell) we used a non-parametric test based on ranks for comparisons.

The type of cell shape did not appear to influence the global anisotropy of the microtubule network. This is interesting as it suggests that different cell types with various shapes do not require differential regulation of the network in order to maintain the anisotropic properties of their CMT arrays (Fig 4A).

**Fig 4.**
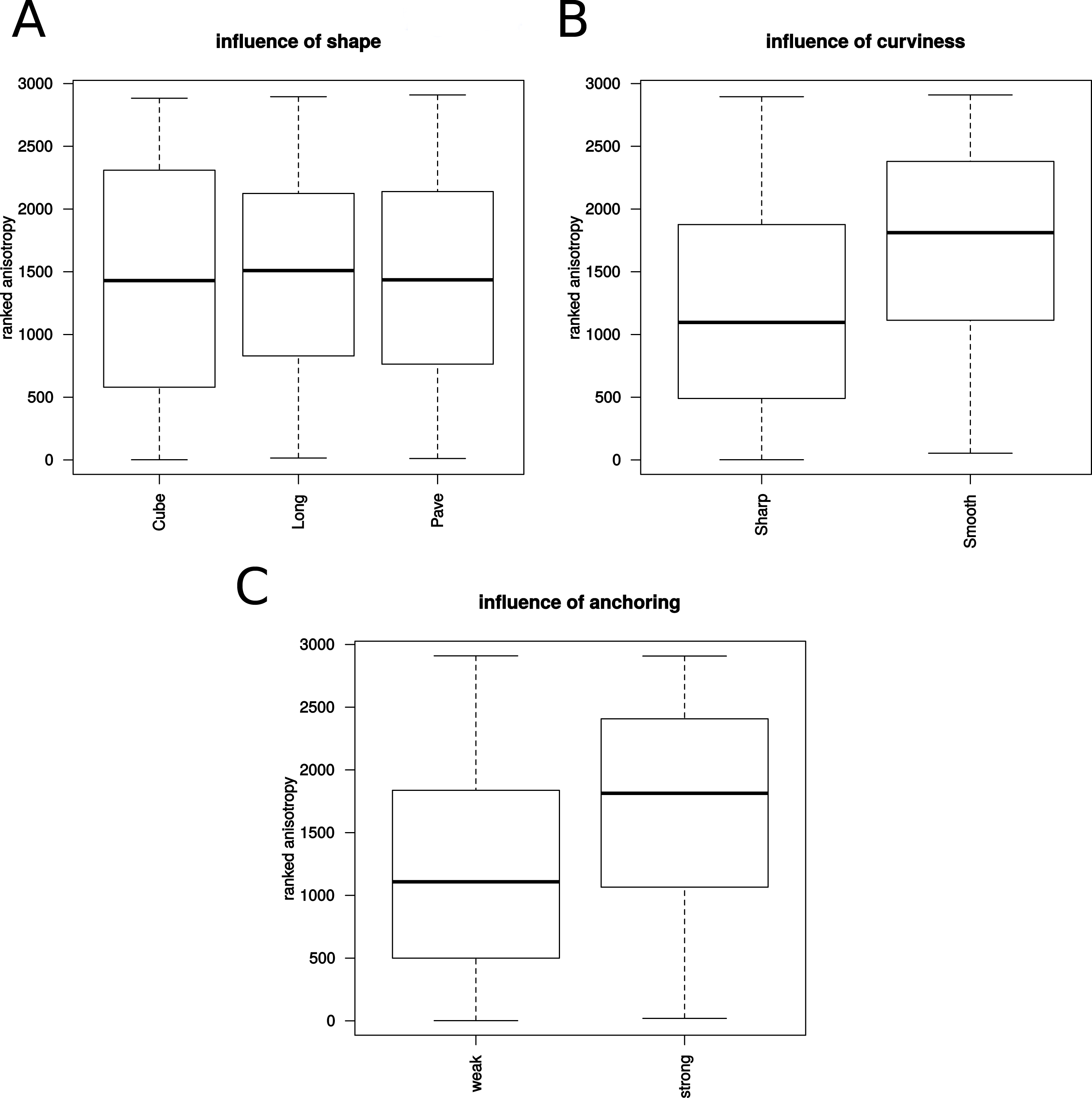
Anisotropy of the network. Influence of shape on (A) anisotropy of the network, (B) of curviness of the walls, and (C) anchoring strength. Anisotropy values are ranked in (A-C).

However, local curvature significantly increased the anisotropy of the microtubule network, by 10% (p<0.001) (Fig 4B). Low convex curvatures that characterize flat cell faces generate less anisotropy in the arrays than higher curvatures. This effect was reduced in the case of elongated cells.

### The average microtubule orientation is strongly influenced by cell shape

In order to determine how the shape of the cell influences the global orientation of the network, we measured the distribution of orientations along two faces of the cells. We chose the top and bottom faces for all shapes, using the sharp squared cell as a reference for determining the labeling of faces. Top and bottom faces are the two larger faces of the cell in our simulations. An angle of 0° corresponds to the long axis in the case of the elongated cell. Angles of 90° or −90° correspond to directions perpendicular to that axis.

First, we observed that in case of square shaped faces, most of the microtubules align along the cell face diagonal, i.e. the longest path (Fig 5). This occurred whatever the anchoring strength and the shape of faces at the side. Second, we observed a strong correlation between the long axis of the cell and microtubule orientation, and this correlation was higher for the elongated cell. Indeed, the microtubule distribution was centered around an angle of 0. This effect is higher in the case of strong anchoring, whereas in the case of weak anchoring, secondary peaks show that the diagonals are also overrepresented. These simulation results indicate that the microtubule network is able to read the longest axis of the cell and orient toward that axis, by default. Interestingly, cortical microtubules become longitudinal in hypocotyl cells when growth stops [5], [38], [44],], suggesting that they may adopt their configuration “by default” in that situation.

**Fig 5.**
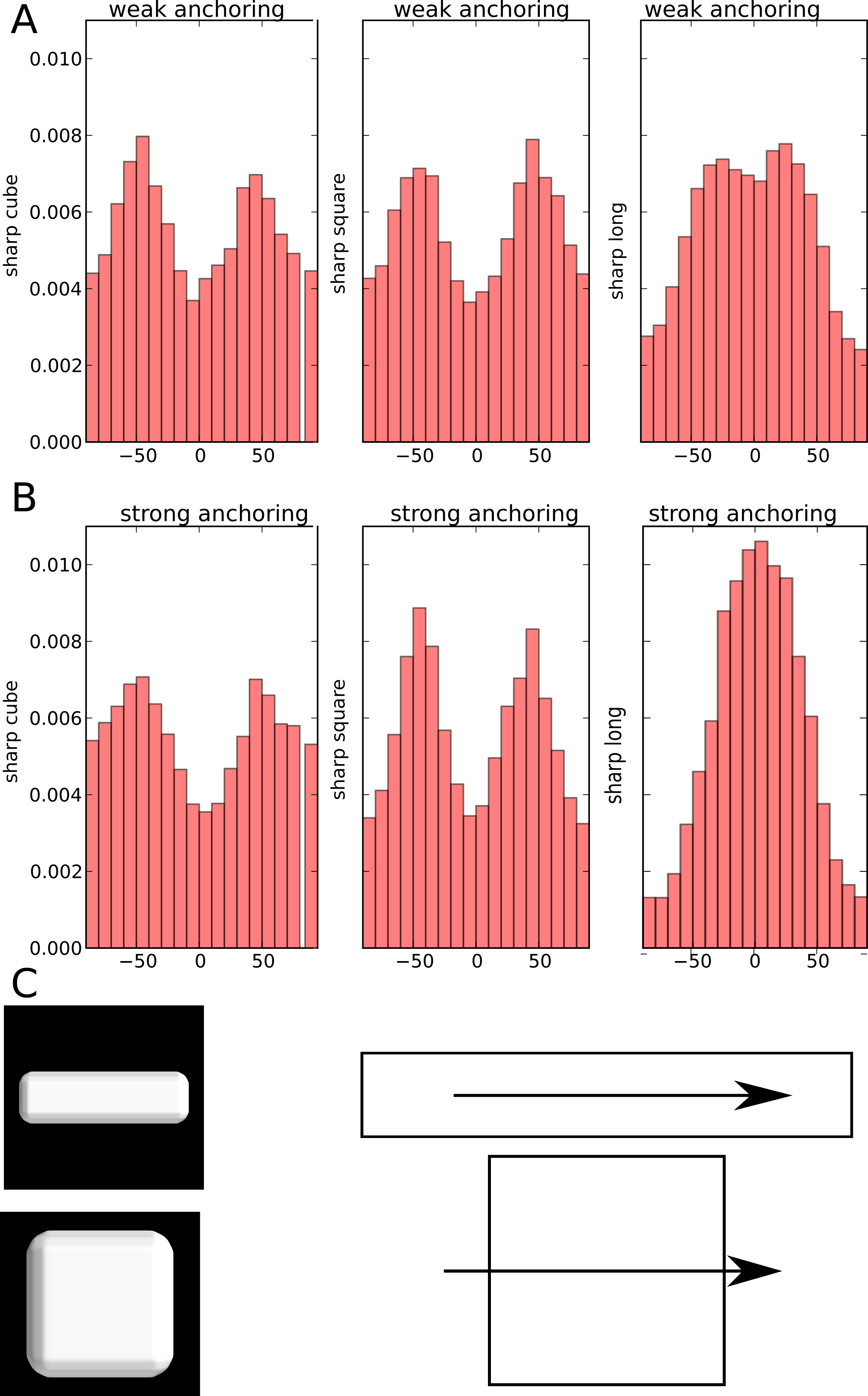
Orientation of the network. Distribution of the orientation of unit vectors (local orientation of microtubules) located close to the top and bottom faces. Top and bottom faces refer to the largest faces of the squared cell represented in C, and to the largest surfaces in the others. The reference angle 0 corresponds to the longest axis of the elongated cell. (A,B) Distribution of orientations for sharp cube, sharp square, sharp long cell, as labeled in Fig 1. (A) Weak anchoring and (B) strong anchoring. (C) Arrow showing the reference for angle measurement for the elongated shape (top) and for the square or the cubic shape (bottom). The shapes are not at the same scale.

### Small external directional bias is sufficient to affect the orientation of the whole network

Next, we investigated the robustness of microtubule network behavior. Microtubule arrays entirely reorganize during cell division [39]. Light and hormones can also completely reorient the microtubule network within minutes [40], [15], [16], suggesting that the constraints on microtubules cannot be too strong to allow such rapid reorganization *in vivo*. We tested whether our model provides such adaptability, using the case of external, directional cues (Fig 6).

**Fig 6.**
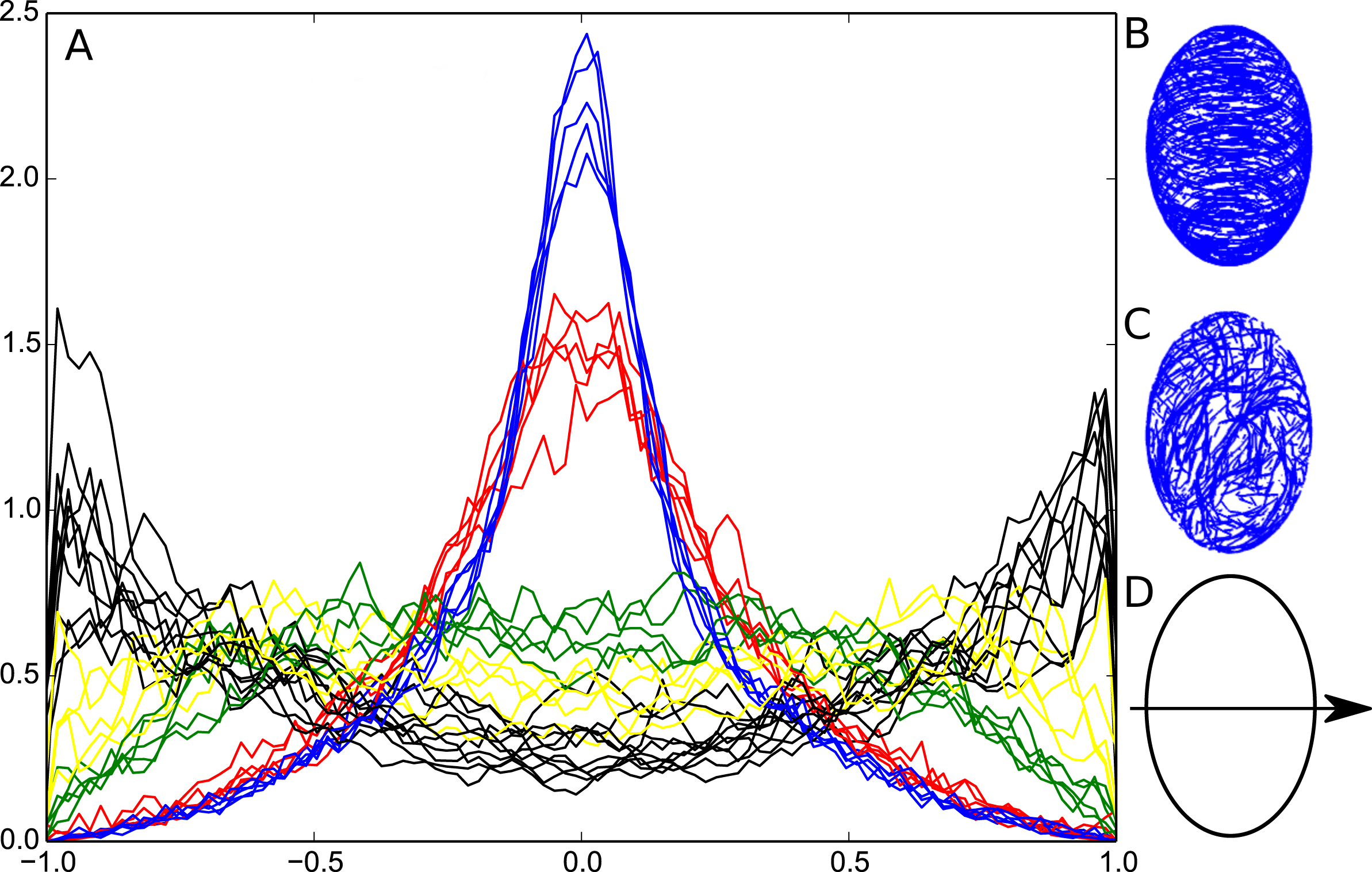
Influence of directional cues on the global orientation of the network. (A) Histogram of the scalar product between the unit vectors (defining microtubules locally) and the main axis of the cell. Black: no directional cue. Colors: directional cue with influence on the direction. The relative weight of the cue is: 0.1% (yellow), 0.2% (green), 1% (red), 2% (blue). (B) Simulation result for a cue weight of 2%. (C) Simulation result with no directional cue. (D) Orientation of the axis of measurement.

We investigated the effect of an external cue, assuming that, as microtubules grow in the cell, their direction is biased toward the cue, with a specific weight. We used an ellipsoidal cell in which the membrane exhibits a directional circumferential cue perpendicular to the main axis of the cell. In the absence of such cue, microtubules are on average parallel to the main axis of the cell. Strikingly, our simulations indicate that even a very low bias (~0.1%) could disrupt the main orientation of the network. When the weight reaches 1% or more, microtubules massively reorient toward the direction of the cue.

These results indicate that, despite an apparent robust organization, the microtubule network remains extremely sensitive to directional cues. As such, it is capable of reading slight directional cues and generating a strong polarity toward that direction. This ability to read directional cues is probably linked to the self-enhancement of microtubule orientation through their interactions: As more microtubules orient toward a direction, they prevent growth of microtubules in the perpendicular direction.

Our simulations indicate that the network should exhibit two behaviors concerning its polarity. When no external directional cue is present, the network orients toward the main axis of the cell, and generates a polarity that is a direct reading of the global shape. When a directional cue is present, the network reorients so as to emphasize the direction of the cue.

## Discussion

### The default state of the microtubule network

The impact of external cues on the microtubule network is well characterized. However, because real cells and tissues are never really devoid of external cues, the behavior of the microtubule network by default in a plant cell remains an open question. Our work provides some clues to address this question, taking into account the 3D shape of the cell and with the most minimal set of parameters for microtubule behavior. We found that microtubule directional persistence largely determines the subcellular localization and orientation of the microtubule network in various cell shapes. We also identified parameters that seemed relatively insensitive to cell shape, such as microtubule length or number. Last, despite the constraints of microtubule stiffness and growth mode on the organization of the microtubule network, we found that such parameters allow high sensitivity to directional cues, even when such cues go against the default orientation. Altogether, this provides a conceptual framework to dissect the exact contribution of microtubule regulators to the microtubule network organization, in relation to 3D cell shape.

### Cortical localization is an emerging property of microtubule stiffness and growth mode in convex cells

Based on TEM images where microtubules are often seen very close to the plasma membrane, it is assumed that anchoring of microtubules to the plasma membrane is relatively strong. However how this anchoring would be mediated remains unknown. Many biochemical studies have been performed to extract proteins that would link the plasma membrane to cortical microtubules, and so far the only published candidate is a phospholipase [41], for which no follow-up results have been obtained to our knowledge. Other links have been put forward, such as CLASP [31] or CELLULOSE SYNTHASE INTERACTIVE PROTEIN 1 (CSI1) [36], [42], but they might represent rather indirect regulators of the link between microtubules and plasma membrane. Does this mean that microtubule could be cortical without any anchoring module? Our results suggest that in most of the measured cell shapes, microtubules do not need to be strongly connected to the membrane to remain cortical. This prediction implies that modulating anchoring would affect self-organizing properties tangentially to the cell surface, rather than modulating the density of microtubules inside the cell. This would allow molecular regulators to modify the microtubule organization without directly affecting the rate of cellulose deposition.

### A high curvature increases anisotropy

In our simulations, we observed differences in anisotropy originating from the sharpness of face-face contacts and curvature of the cell. Cells that are more curved exhibit increased array anisotropy compared to flatter ones. This result is interesting, as epidermal cells possess faces with very different curvatures. In the L1 layer of the shoot apical meristem for example, the outermost wall has a stronger curvature than all the walls separating the cell from its neighbors. Pavement cells also display various local curvature values. Based on our results, higher anisotropy would be produced in L1 cells without specific regulation. Such results are difficult to account for in models with 2D surfaces embedded in 3D [31].

Similarly, changes in the curvature of the epidermal wall can occur through changes in internal pressure, which in return influence the anisotropy of the network [43]. An increase in the microtubule organization could be a result of an increased pressure, that would increase the curvature of the epidermal cell. Further experimental work is required to investigate the role of curvature on microtubule behavior and its relation to mechanical stress in the epidermis.

### Influence of shape on the global directionality of the network

Our simulations show that the cell aspect ratio has an impact on the global orientation of the network. The predicted default behavior of microtubules is their alignment parallel to the long axis of a cell, due to the directional persistence of microtubules associated with their bending stiffness. This is in agreement with previous models with microtubules living on 2D surfaces embedded in 3D [34], [35], where the default orientation is longitudinal for long cylinders.

This default state was observed with microtubules polymerizing in vitro inside elongated chambers [34]. In slowly growing cells of the hypocotyl, microtubules are oriented along the long axis of the cell, whereas microtubules are circumferential in rapidly elongating cells [38], [44]. Our model suggests that directional cues are needed to avoid this default orientation is growing cells.

At the boundary between the shoot apical meristem and the primordia, cell division leads to cell shapes that are elongated along the axis of the boundary; our model predicts that microtubules will be oriented along the same direction by default, amplifying their response to mechanical stress [8].

### A weak directional bias is sufficient to change the orientation of the microtubule network

In this study, we show that the microtubule network is oriented by default along the longest axis of the cell. However, microtubules in plants often show supracellular orientation, independently of cell shape, a behavior that has been ascribed to tissue-level signals, notably mechanical stress [8]. Moreover, It has been demonstrated that inside the cell, microtubules orientation relates to polarity markers such as proteins from the PIN FORMED and RHO OF PLANTS families [45], [46]. Simulations have assessed how localized membrane heterogeneity could result in a biased orientation of the microtubule network [31], [30]. In this study, we show that a weak directional cue influencing microtubule growth rapidly modifies the orientation of the network towards the direction of the cue. As such, the network behaves as a perception tool translating an external directional information into a structural polarity inside the cell. The coexistence of a default orientation and a strong ability to reorient could shed a new light on orientation changes in cells. Changes in microtubule orientation need not be always related to specific regulation but my also be related to the arrest of signals and the return to the default state. This could be occurring in the shift from transverse to longitudinal orientation in hypocotyls responding to light or hormones [40], [15], [16].

### Perspectives

The shape of the cell has little influence on mean length, number of microtubules or bundles proportions. In addition, anisotropy of the network is not highly correlated to changes in global cell shape. This prediction of a robust network suggests that plant cells do not need specific regulations to compensate for their great variations in shapes. Accordingly, the microtubule network appears as a good polarity system, with a default orientation and a high sensitivity to directional cues. It was recently shown that, for global polarity to emerge in a tissue, the only requirement is the existence of internal cellular polarity [47]. In this work we show that the microtubule network is adapted to such requirement.

Overall, microtubules and associated proteins form a complex self-organizing system that is difficult to comprehend without resorting to models. The results obtained here demonstrate that our three-dimensional model provides a framework to test hypotheses on the regulation of the microtubule cytoskeleton in plant cells.

The model given here is a beginning to a more complete analysis. We have not yet incorporated microtubule severing [40], [28], microtubule branching [11], and the possible effects of connections between cortical microtubules and cellulose fibrils outside of the cell as mediated by the cellulose synthase complex [48]. Severing in particular has been shown to be key to microtubule reorientation [28] following mechanical signals [49]. We also have not included limiting levels of tubulin [35], which could affect overall microtubule number. Altogether, we expect our model to help progress in understanding how microtubule self-organization integrates three-dimensional cell shape and directional cues and how microtubule-associated proteins modulate this integration.

## Methods

The dynamical microtubule model was implemented in C++. Simulations were performed on Intel/AMD desktop computers running Debian and Ubuntu operating systems.

### The microtubule network

Microtubules were coded as 3D multi-segment vectors of constant length. A ring of tubulin of the height of a dimer is represented as a unit vector in the simulation. Microtubule growth in the model occurs by adding one vector element at the plus end of the microtubule, at a position in the direction pointed to by the last vector. In plants, microtubules are considered static, their growth and shrinkage are the result of treadmilling processes. To code for microtubule directional persistence (which relates physically to bending stiffness), the direction at which a new vector is added to an existing microtubule changes by a small random amount. Microtubule shrinkage occurs at the minus end by removing the first vector from the list.

### Time and space correspondences

Typically, a cell has a width of several micrometers, and we take the unit of length as 8nm, the height of a ring of tubulin. Considering a measured speed of growth at plus ends of 3-5 µm/min [50], a simulation time step is approximately 0.1 to 0.2 s. A typical simulation of 10000 time steps thus represents 16 to 27 min of real time.

### Nucleation and minus end rules

The microtubules are nucleated close to the cell surface at a rate of 1 to 20 microtubules/s. Once nucleated, the microtubules do not immediately shrink. At each of the time steps that follow nucleation, a microtubule has a probability *n*_s_ to begin shrinkage.

### Plus-end growth

Without any interactions with the membrane or with neighbouring microtubules, the new direction of growth is that of the previous vector (tubulin), modified by a random direction *r*_d_, typically deviating less than 10° from the previous direction. This corresponds to a persistence length of a few micrometers. In the presence of directional cue, the new vector is computed as the weighted sum of its unbiased direction and the direction of the cue vector. When a microtubule reaches the plasma membrane, it follows one of two behaviors depending on the interaction strength with the membrane. a) strong anchoring: after interaction, the microtubule grows along to the surface with further new vectors added tangentially to the surface, or b) weak anchoring: the microtubule grows as in the case of free space, except that it cannot go out of the cell: if the new vector is outside the cell then the vector is projected on the plane tangential to the surface of the cell; due to the noise on vector orientation, the microtubule may leave cell surface.

When a plus end is closer than 25 nm to another microtubule, two situations can occur: a) if the angle between the microtubule and the encountered one is smaller than a threshold α, the microtubule begins zippering, and its direction becomes that of the previously existing microtubule; b) if the angle is larger than α, the microtubule begins to shrink at the plus end (catastrophe event) or stalls according to the encounter rule. α was set to 40° [20]

**Table 1:** Main parameters for microtubule dynamics

| Parameter | Nucleation | Shrinkage | Random | Encounter | Anchoring | Plus end |
| --- | --- | --- | --- | --- | --- | --- |
|  | frequency | probability | direction | angle | strength | encounter rule |
| Label | $n_p$ | $n_s$ | $r_d$ | $\alpha$ | | |
| Default value | 2 nucleations<br>par time step | 0.001 per nucleation<br>site per time step | <0.03 | $40^\circ$ | Strong or<br>weak | Catastrophe or<br>stall |

### Cell shape and interaction with the membrane

The cell contour is described with a triangular mesh. Each vertex possesses the information of the vector normal to the surface, which is used during the simulation to calculate a local approximation of the tangential plane. It is possible to add information at the vertex level that can be read during the simulation and serve as extrinsic input. Inputs can be scalars or tensors. The distance to the membrane is calculated using the nearest point on the surface. At this point, the membrane is approximated by the plane perpendicular to the normal of the mesh. The distance between a tubulin ring (unit vector) and the membrane is calculated as the shortest distance between the endpoint of the vector and this plane. A collection of standard cell shapes was generated for our simulations (Fig. 1A).

### The simulation process

The simulation progress is made through discrete timesteps. At each timestep new vectors are added and vectors are removed from the simulation space. Collision tests are realized so as to implement the different growth or shrinkage situations. In order to increase the simulation speed, space is divided into subelements and the vectors are identified according to the subelement to which they belong, which diminishes the number of particles involved in the collision test (locality-sensitive hashing [51].

### Visualization

To visualize microtubule density and orientations an image is created using a matrix of resolution rx,ry,rz. rz is larger relative to rx and ry (typically 10 to 1 ratio), which mimics the anisotropic quality of confocal microscope resolution. The simulation space is then screened. When a vector is located inside a cube of the matrix, the value of this cube is incremented by one, and the immediate neighbouring cubes are incremented by a lower number (typically 0.3). At the end of the process, a stack is formed where microtubules appear as blurred intensity signals. One can either visualize each subimage from the stack by moving along the z axis, or create a projection that sums the matrix along the z axis.

### Quantification of anisotropy of microtubule arrays

The space is subdivided into cubes of arbitrary size, typically segmenting the structure into circa 27 cubes. Segmenting the structure into 216 smaller pieces gives similar results, with globally higher anisotropy values. All tubulin ring directions are extracted as a 3xN matrix: D. The three eigenvalues λ_1_, λ_2_, λ_3_ are obtained through the diagonalisation of the 3x3 matrix 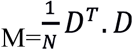

Anisotropy, A, of the microtubule array in each cube of space is quantified by

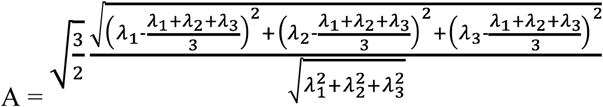

Anisotropies are then averaged over the whole cell.

## Acknowledgments

We thank the members of RDP and Sainsbury laboratories for discussions and support.

## Bibliography

1. Ledbetter MC. A “MICROTUBULE” IN PLANT CELL FINE STRUCTURE. J Cell Biol. 1963;19: 239–250. doi:10.1083/jcb.19.1.239

2. Brandizzi F, Wasteneys GO. Cytoskeleton-dependent endomembrane organization in plant cells: an emerging role for microtubules. Plant J. 2013;75: 339–49. doi:10.1111/tpj.12227

3. Baskin, Meekes, Liang, Sharp. Regulation of growth anisotropy in well-watered and water-stressed maize roots. II. Role Of cortical microtubules and cellulose microfibrils. Plant Physiol. 1999;119: 681–92. Available: http://www.plantphysiol.org/content/119/2/681.short

4. Paredez AR, Somerville CR, Ehrhardt DW. Visualization of cellulose synthase demonstrates functional association with microtubules. Science. 2006;312: 1491–5. doi:10.1126/science.1126551

5. Baskin TI. Anisotropic expansion of the plant cell wall. Annu Rev Cell Dev Biol. 2005;21: 203–22. doi:10.1146/annurev.cellbio.20.082503.103053

6. Bringmann M, Landrein B, Schudoma C, Hamant O, Hauser M-T, Persson S. Cracking the elusive alignment hypothesis: the microtubule-cellulose synthase nexus unraveled. Trends Plant Sci. Elsevier Ltd; 2012;17: 666–74. doi:10.1016/j.tplants.2012.06.003

7. Buschmann H, Lloyd CW. Arabidopsis mutants and the network of microtubule-associated functions. Mol Plant. 2008;1: 888–898. doi:10.1093/mp/ssn060

8. Hamant O, Heisler MG, Jönsson,H, Krupinski P, Uyttewaal M, Bokov P, et al. Developmental patterning by mechanical signals in Arabidopsis. Science. 2008;322: 1650–5. doi:10.1126/science.1165594

9. Sampathkumar A, Yan A, Krupinski P, Meyerowitz EM. Physical forces regulate plant development and morphogenesis. Curr Biol. 2014;24: R475–R483. doi:10.1016/j.cub.2014.03.014

10. Fishel EA, Dixit R. Role of nucleation in cortical microtubule array organization: variations on a theme. Plant J. 2013;75: 270–277. doi:10.1111/tpj.12166

11. Chan J, Sambade A, Calder G, Lloyd C. Arabidopsis Cortical Microtubules Are Initiated along, as Well as Branching from, Existing Microtubules. PLANT CELL ONLINE. 2009;21: 2298–2306. doi:10.1105/tpc.109.069716

12. Ambrose C, Wasteneys GO. Microtubule initiation from the nuclear surface controls cortical microtubule growth polarity and orientation in Arabidopsis thaliana. Plant Cell Physiol. 2014;55: 1636–1645. doi:10.1093/pcp/pcu094

13. Mitchison T, Kirschner M. Dynamic instability of microtubule growth. Nature. 1984;312: 237–42. doi:10.1038/312237a0

14. Shaw SL, Kamyar R, Ehrhardt DW. Sustained microtubule treadmilling in Arabidopsis cortical arrays. Science. 2003;300: 1715–8. doi:10.1126/science.1083529

15. Sambade A, Pratap A, Buschmann H, Morris RJ, Lloyd C. The influence of light on microtubule dynamics and alignment in the Arabidopsis hypocotyl. Plant Cell. 2012;24: 192–201. doi:10.1105/tpc.111.093849

16. Vineyard L, Elliott A, Dhingra S, Lucas JR, Shaw SL. Progressive transverse microtubule array organization in hormone-induced Arabidopsis hypocotyl cells. Plant Cell. 2013;25: 662–676. doi:10.1105/tpc.112.107326

17. Zhang Q, Fishel E, Bertroche T, Dixit R. Erratum: Microtubule severing at crossover sites by katanin generates ordered cortical microtubule arrays in arabidopsis (Current Biology (2013) 23 (2191-2195)). Current Biology. 2014. p. 917. doi:10.1016/j.cub.2014.03.057

18. Wasteneys GO, Ambrose JC. Spatial organization of plant cortical microtubules: close encounters of the 2D kind. Trends Cell Biol. 2009;19: 62–71. doi:10.1016/j.tcb.2008.11.004

19. Allard JF, Ambrose JC, Wasteneys GO, Cytrynbaum EN. A mechanochemical model explains interactions between cortical microtubules in plants. Biophys J. Biophysical Society; 2010;99: 1082–1090. doi:10.1016/j.bpj.2010.05.037

20. Dixit R, Cyr R. Encounters between dynamic cortical microtubules promote ordering of the cortical array through angle-dependent modifications of microtubule behavior. Plant Cell. 2004;16: 3274–84. doi:10.1105/tpc.104.026930

21. Stoppin-Mellet V, Gaillard J, Vantard M. Katanin’s severing activity favors bundling of cortical microtubules in plants. Plant J. 2006;46: 1009–1017. doi:10.1111/j.1365-313X.2006.02761.x

22. Baulin VA, Marques CM, Thalmann F. Collision induced spatial organization of microtubules. Biophys Chem. 2007;128: 231–44. doi:10.1016/j.bpc.2007.04.009

23. Shi X -q. X, Ma Y. Understanding phase behavior of plant cell cortex microtubule organization. Proc Natl Acad Sci. 2010;107: 11709–11714. doi:10.1073/pnas.1007138107

24. Hawkins RJ, Tindemans SH, Mulder BM. Model for the orientational ordering of the plant microtubule cortical array. Phys Rev E Stat Nonlin Soft Matter Phys. 2010;82: 11911. doi:10.1103/PhysRevE.82.011911

25. Tindemans SH, Hawkins RJ, Mulder BM. Survival of the aligned: Ordering of the plant cortical microtubule array. Phys Rev Lett. 2010;104: 58103. doi:10.1103/PhysRevLett.104.058103

26. Allard JF, Wasteneys GO, Cytrynbaum EN. Mechanisms of self-organization of cortical microtubules in plants revealed by computational simulations. Mol Biol Cell. 2010;21: 278–286. doi:10.1091/mbc.E09-07-0579

27. Deinum EE, Tindemans SH, Mulder BM. Taking directions: the role of microtubule-bound nucleation in the self-organization of the plant cortical array. Phys Biol. 2011;8: 56002. doi:10.1088/1478-3975/8/5/056002

28. Deinum EE, Tindemans SH, Lindeboom JJ, Mulder BM. How selective severing by katanin promotes order in the plant cortical microtubule array. Proc Natl Acad Sci U S A. 2017;114: 6942–6947. doi:10.1073/pnas.1702650114

29. Lindeboom JJ, Lioutas A, Deinum EE, Tindemans SH, Ehrhardt DW, Emons AMC, et al. Cortical microtubule arrays are initiated from a nonrandom prepattern driven by atypical microtubule initiation. Plant Physiol. 2013;161: 1189–201. doi:10.1104/pp.112.204057

30. Eren EC, Dixit R, Gautam N. A three-dimensional computer simulation model reveals the mechanisms for self-organization of plant cortical microtubules into oblique arrays. Mol Biol Cell. 2010;21: 2674–84. doi:10.1091/mbc.E10-02-0136

31. Ambrose C, Allard JF, Cytrynbaum EN, Wasteneys GO. A CLASP-modulated cell edge barrier mechanism drives cell-wide cortical microtubule organization in Arabidopsis. Nat Commun. Nature Publishing Group; 2011;2: 430. doi:10.1038/ncomms1444

32. Buxton GA, Siedlak SL, Perry G, Smith MA. Mathematical modeling of microtubule dynamics: insights into physiology and disease. Prog Neurobiol. Elsevier Ltd; 2010;92: 478–83. doi:10.1016/j.pneurobio.2010.08.003

33. Nedelec F, Foethke D. Collective Langevin dynamics of flexible cytoskeletal fibers. New J Phys. 2007;9. doi:10.1088/1367-2630/9/11/427

34. Cosentino Lagomarsino M, Tanase C, Vos JW, Emons AMC, Mulder BM, Dogterom M. Microtubule organization in three-dimensional confined geometries: evaluating the role of elasticity through a combined in vitro and modeling approach. Biophys J. Elsevier; 2007;92: 1046–57. doi:10.1529/biophysj.105.076893

35. Tindemans SH, Deinum EE, Lindeboom JJ, Mulder BM. Efficient event-driven simulations shed new light on microtubule organization in the plant cortical array. Front Phys. Frontiers; 2014;2: 1–15. doi:10.3389/fphy.2014.00019

36. Li S, Lei L, Somerville CR, Gu Y. Cellulose synthase interactive protein 1 (CSI1) links microtubules and cellulose synthase complexes. Proc Natl Acad Sci U S A. 2012;109: 185–90. doi:10.1073/pnas.1118560109

37. Mei Y, Gao H-B, Yuan M, Xue H-W. The Arabidopsis AR CP protein, CSI1, which is required for microtubule stability, is necessary for root and anther development. Plant Cell. 2012;24: 1066–80. doi:10.1105/tpc.111.095059

38. Crowell EF, Timpano H, Desprez T, Franssen-Verheijen T, Emons A-M, Höfte,H, et al. Differential regulation of cellulose orientation at the inner and outer face of epidermal cells in the Arabidopsis hypocotyl. Plant Cell. 2011;23: 2592–605. doi:10.1105/tpc.111.087338

39. Müller S. Universal rules for division plane selection in plants. Protoplasma. 2012;249: 239–53. doi:10.1007/s00709-011-0289-y

40. Lindeboom JJ, Nakamura M, Hibbel A, Shundyak K, Gutierrez R, Ketelaar T, et al. A mechanism for reorientation of cortical microtubule arrays driven by microtubule severing. Science. 2013;342: 1245533. doi:10.1126/science.1245533

41. Gardiner JC, Harper JD, Weerakoon ND, Collings D a, Ritchie S, Gilroy S, et al. A 90-kD phospholipase D from tobacco binds to microtubules and the plasma membrane. Plant Cell. 2001;13: 2143–58. doi:10.1105/TPC.010114

42. Bringmann M, Li E, Sampathkumar A, Kocabek T, Hauser M-T, Persson S. POM-POM2/CELLULOSE SYNTHASE INTERACTING1 Is Essential for the Functional Association of Cellulose Synthase and Microtubules in Arabidopsis. Plant Cell. 2012;24: 163–177. doi:10.1105/tpc.111.093575

43. Liu Q, Qiao F, Ismail A, Chang X, Nick P. The plant cytoskeleton controls regulatory volume increase. Biochim Biophys Acta - Biomembr. Elsevier B.V.; 2013;1828: 2111–2120. doi:10.1016/j.bbamem.2013.04.027

44. Chan J, Eder M, Crowell EF, Hampson J, Calder G, Lloyd C. Microtubules and CESA tracks at the inner epidermal wall align independently of those on the outer wall of light-grown Arabidopsis hypocotyls. J Cell Sci. 2011;124: 1088–1094. doi:10.1242/jcs.086702

45. Heisler MG, Hamant O, Krupinski P, Uyttewaal M, Ohno C, Jönsson,H, et al. Alignment between PIN1 Polarity and Microtubule Orientation in the Shoot Apical Meristem Reveals a Tight Coupling between Morphogenesis and Auxin Transport. Leyser O, editor. PLoS Biol. 2010;8: e1000516. doi:10.1371/journal.pbio.1000516

46. Fu Y, Xu T, Zhu L, Wen M, Yang Z. A ROP GTPase Signaling Pathway Controls Cortical Microtubule Ordering and Cell Expansion in Arabidopsis. Curr Biol. 2009;19: 1827–1832. doi:10.1016/j.cub.2009.08.052

47. Abley K, De Reuille PB, Strutt D, Bangham A, Prusinkiewicz P, Maree AFM, et al. An intracellular partitioning-based framework for tissue cell polarity in plants and animals. Development. 2013;140: 2061–2074. doi:10.1242/dev.062984

48. Paradez A, Wright A, Ehrhardt DW. Microtubule cortical array organization and plant cell morphogenesis. Curr Opin Plant Biol. 2006;9: 571–578. doi:10.1016/j.pbi.2006.09.005

49. Uyttewaal M, Burian A, Alim K, Landrein B, Borowska-Wykrt D, Dedieu A, et al. Mechanical stress acts via Katanin to amplify differences in growth rate between adjacent cells in Arabidopsis. Cell. 2012;149: 439–451. doi:10.1016/j.cell.2012.02.048

50. Ehrhardt DW, Shaw SL. MICROTUBULE DYNAMICS AND ORGANIZATION IN THE PLANT CORTICAL ARRAY. Annu Rev Plant Biol. 2006;57: 859–875. doi:10.1146/annurev.arplant.57.032905.105329

51. Gionis A, Indyk P, Motwani R. Similarity Search in High Dimensions via Hashing. VLDB’99 Proc 25th Int Conf Very Large Data Bases. 1999;99: 518–529. doi:10.1.1.41.4809

